# From Wiring to Firing: Collapse of embryonic identities and emergence of functional diversity during motor neuron maturation

**DOI:** 10.1101/2025.08.12.669897

**Authors:** Yijia Chen, Hsuan-Ming Chi, Aileen Tian, Alison Miller, Tulsi Patel

## Abstract

Neurons born in the embryo undergo a protracted process of maturation during which time they project axons to their specific targets, integrate into circuits, refine synapses, and acquire unique electrophysiological properties. The molecular strategies that individual neuron types deploy to complete this complex process remain poorly understood. In this work, we used single nucleus multiome sequencing (RNA-seq and ATAC-seq) to track the transition from specification to functional maturation in mouse skeletal motor neurons (SMNs). Our data show that individual SMNs undergo significant transcriptional changes as they mature, but more strikingly, we find that diversity within SMNs fluctuates dramatically as the functional needs of these cells change over time. At embryonic day E15.5, when motor axons are innervating their specific muscle targets, SMNs can be subdivided into dozens of transcriptional subclusters. These embryonic subclusters represent known motor columns and pools, which utilize column- and pool-specific genes to innervate unique muscle targets. About a week later, at postnatal day 3 (P3) many column- and pool-specific genes are downregulated or become broadly expressed and SMNs coalesce on the molecular level into a more homogenous state. These neurons then undergo a second round of diversification during the first two weeks of postnatal life (P3-P13), acquiring gene expression patterns that divide them into the functionally distinct alpha, gamma, and type3 subtypes found in adults. The fluctuations in SMN diversity go hand-in-hand with changes in accessible chromatin regions and transcription factor (TF) expression. Differential ATAC-seq peaks that define embryonic diversity are lost over time while new peaks that control expression in adult subtypes are gained. TFs that are known to regulate embryonic diversity are also downregulated over time, as a separate set of TFs that likely regulate adult subtype identities are upregulated. Our work uncovers a novel maturation trajectory for postmitotic neurons where extensive spatial diversity is first acquired in the embryo to ensure proper circuit wiring; this diversity is then lost as maturing neurons re-diversify into functional identities required for proper circuit firing in postnatal life. Therefore, all aspects of a neuron’s identity – its morphology, circuitry, and electrophysiologically – may not be fully described by its gene expression program at adulthood, but instead is a culmination of transcriptional events that occur throughout its specification and maturation trajectory as the functional needs of the cells evolve.

## Introduction

Adult neurons are remarkably diverse in their morphology, circuitry, and physiology. This diversity is initiated during embryonic development, when progenitors and newly born postmitotic neurons activate unique gene expression programs in response to spatial signaling cues^1,2^. Nascent neurons then undergo a prolonged period of maturation as they extend projections, integrate into circuits, and refine synapses^3^. This process can take years in humans and weeks in mice and coincides with the acquisition of mature behaviors in organisms^4^. Recent genomics studies have demonstrated that neuronal maturation is also accompanied by neuron-type specific transcriptional changes, such that gene expression programs of adult neurons are remarkably different from embryonic neurons^5–9^. Decades of neurodevelopmental studies have elucidated mechanisms of embryonic diversity, but only recently – with the advances in single cell sequencing methods – have we begun to understand how this diversity changes as neurons mature into their adult forms.

Single cell maturation trajectories from embryo to adulthood have now been mapped for a few mammalian neuron types including cortical neurons, hypothalamic neurons, and dorsal root ganglia sensory neurons^6,7,9–13^. These studies demonstrate that while all neuron types show gene expression changes as they mature, their diversification into adult subtypes can follow distinct trajectories. For example, single cell RNA-seq data^6^ shows that sensory neurons in the mouse dorsal root ganglia are fairly homogenous after specification in the embryo, but they diversify into functional subtypes by early postnatal life, and maintain this diversity into adulthood. Similarly, cortical inhibitory and excitatory neurons in humans and mice also become more diverse as they transition from embryonic to postnatal states^9,14,15^. In contrast, inhibitory and excitatory neurons in the preoptic area of the hypothalamus are already highly diverse in the embryo and they maintain the same level of diversity through adulthood^7^. Therefore, different neuron types seem to acquire adult-like diversity at different stages of development and maturation, raising questions about how these trajectories are regulated and why they are divergent between neuron types.

To gain further insight into neuron-type specific transitions from embryonic to adult diversity, here we focus on mouse skeletal motor neurons (SMNs). SMNs reside in the spinal cord and innervate muscles throughout the body to control movement and coordination. All SMNs are born during embryonic development, but their circuits, synapses, and electrophysiological properties continue to mature during postnatal life as animals progressively control posture, crawl, walk, and run^16–18^. SMNs make an intriguing cell type for this study because the transcriptional diversity described in functionally mature adult SMNs^19,20^ is striking different from transcriptional diversity described in embryonic SMNs^21^, but it remains unclear how and when the transition from embryonic diversity to adult diversity takes place^22^.

In the embryo, all post-mitotic SMNs are specified by the action of signaling factors retinoic acid and sonic hedgehog, which lead to activation of selector TFs Isl1 and Lhx3^23–25^. Within ∼3-5 days of specification, post-mitotic SMNs diversify into molecularly-distinct columnar and pool identities along the length of the spinal cord through the action of rostro-caudal signaling factors, patterned expression of Hox genes and other TFs, and signals from specific muscle targets^1,26,27^. This diversification leads to the formation of five discrete SMN clusters called motor columns which further divide into dozens of distinct motor pools^1,21,28^. All SMNs in specific motor columns innervate spatially distinct groups of muscles – for example SMNs in the Lateral Motor Column innervate limb muscles while SMNs in the Medial Motor Column innervate epaxial muscles – and all SMNs in a specific motor pool innervate a single muscle. This extensive spatial diversity in embryonic SMNs therefore ensures the correct wiring of SMN to muscle circuits.

Surprisingly, in recent single nucleus RNA-seq studies, adult SMNs were shown to subdivide into just three main transcriptional identities, as opposed to the dozens of column and pool identities that exist in the embryo^19,20,29^. These three transcriptional identities likely correspond to three functionally distinct adult SMN subtypes – alpha, gamma, and beta – that were first characterized in the 1950s and 1960s based on their muscle innervation patterns. Alpha SMNs innervate extrafusal muscle fibers in each muscle, which control muscle contraction; gamma SMNs innervate intrafusal muscle spindles which control muscle stretch; and beta SMNs innervate both extrafusal fibers and muscle spindle^30^. While very little is understood about beta SMNs (and we therefore refer to them as ‘type3’^20^), alpha and gamma SMNs are known to be morphologically and electrophysiologically distinct. Importantly, these subtypes also show differential susceptibility to diseases like Amyotrophic Lateral Sclerosis – alpha SMNs degenerate while gamma SMNs seem to be spared^31,32^. Although these functionally distinct and disease relevant adult SMN subtypes were described over 60 years ago, their developmental trajectory remains unknown. It is unclear when and how these subtypes are specified and how three functional subtypes emerge from dozens of spatially defined embryonic identities.

To uncover when and how the transition from embryonic diversity to adult diversity takes place, we performed paired single nucleus RNA-seq and ATAC-seq (snMultiome) at six time points encompassing embryonic (E15.5), early postnatal (P3 and P7), juvenile (P13 and P21) and adult (P56) stages. Strikingly, we found that SMN diversity fluctuates significantly during maturation as the functional needs of these cells change. At E15.5, we see SMNs divide into dozens of distinct pool/column identities that are required for proper muscle targeting. However, after muscle connections are established, these identities are lost during perinatal ages and SMNs become more transcriptionally homogenous. Then, during postnatal maturation, SMNs diversify again into the three functional subtypes found in adults, which allows for correct control of muscle contraction and stretch. We found that this fluctuation in diversity is accompanied by a downregulation of embryonic TFs and open chromatin regions, and a gain of new open chromatin regions and TFs during postnatal maturation, demonstrating that molecular signatures of pool and column identities are largely forgotten by adulthood, and that SMN development occurs in distinct steps. Overall, our work highlights the importance of studying neuronal development across the maturation timescale and demonstrates that the identity of a long living postmitotic cell type can change dramatically over time to meet evolving functional needs.

## Results

### A temporal single nucleus multiome atlas of maturing spinal cord cholinergic neurons

To understand the maturation trajectory of adult SMN subtypes, we performed snMultiome sequencing at six timepoints ranging from embryo to adulthood: E15.5, P3, P7, P13, P21, and P56. Postmitotic SMNs are specified at E9.5 and their axons start reaching muscle targets at E11.5^1^. Between E11.5 and E15.5 a wave of developmentally programmed cell death eliminates ∼50% of all SMNs^33^. We start our single nucleus analysis at E15.5 to track only the SMNs that survive and are maintained during postnatal life. To enrich for SMNs, we sorted GFP+ nuclei from cervical and lumbar enriched regions of spinal cord of Chat-Cre;Sun1-sfGFP-myc mice^34,35^ (**Fig. 1a-1c**). These mice express the nuclear membrane marker Sun1-sfGFP in cholinergic neurons including SMNs^5,20^ (**Fig. 1b**). Nuclei from six to ten mice of both sexes were pooled for multiome sequencing (10X Genomics) at each age. After quality control filtering based on both RNA-seq and ATAC-seq reads (details in Methods), we retained 44,088 total nuclei from all ages. To determine the cellular composition of our sequenced nuclei, we performed UMAP clustering and checked expression of known markers (**Fig 1d**). At all ages, we identified clusters of Chat+, Slc5a7+ cholinergic neurons that express established SMN markers Gfra1, Bcl6, and Ret. In addition, we also identified a clear population of cholinergic preganglionic motor neurons (PGMNs, referred to as VMNs in^19,20^) which express Zeb2 and Fbn2, and cholinergic interneurons that express Pitx2 at all ages (**Supplementary Fig. 1a**). In addition to Chat+ cholinergic neurons, the sequenced neurons also included Slc17a6+ Chat- clusters and Gad1+ Chat- clusters at postnatal ages (**Supplementary Fig. 1a**). This is in agreement with a previous single nucleus RNA-seq study which found a similar distribution of cell types in GFP+ nuclei of Chat-Cre;Sun1-sfGFP adult mouse spinal cords^20^. Our clusters from E15.5 and P3/P7 suggest that Chat is transiently co-expressed with Slc17a6 and Gad1 in the embryo and early postnatal life, leading to Sun1-sfGFP labeling of these neurons that is retained even after Chat expression is lost. Overall, this dataset allows us to track the maturation trajectory of all cholinergic neurons in the spinal cord at the single cell level. In this work we focus on the maturation trajectory of Chat+ SMNs, with some comparisons to Chat+ PGMNs.

**Figure 1:**
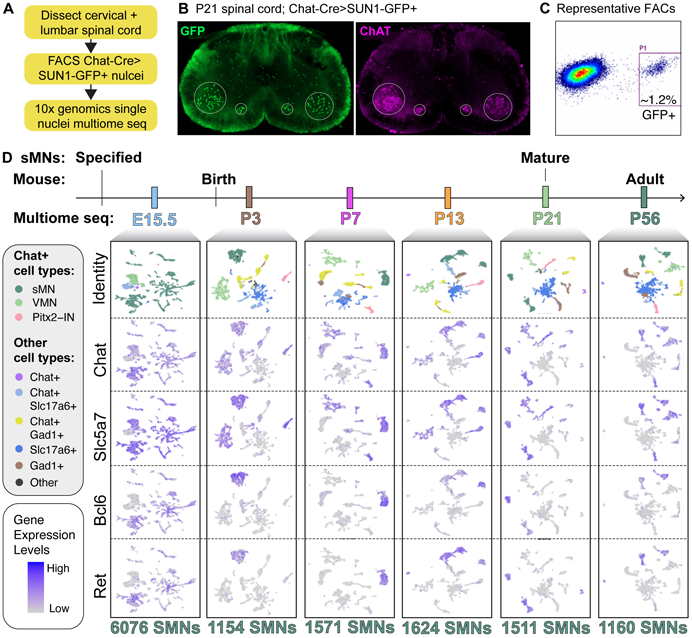
Single nucleus multiome maturation map of spinal cord cholinergic neurons. **A)** Outline of single nucleus multiome experiment. **B)** P21 spinal cord section from Chat-Cre>SUN1-sfGFP-myc mice. SUN1-sfGFP is in green, CHAT protein in magenta. At P21, SUN1-GFP is expressed in neurons that are both positive and negative for CHAT protein, which is consistent with identity of sequenced nuclei in D. **C)** Representative FACs profile of GFP+ nuclei. **D)** UMAP representation of all nuclei that passed quality control at each age. Identity of nuclei is determined by expression of cholinergic genes Chat and Slc5a7, GABAergic gene Gad1, and excitatory gene Slc17a6, as well as cell type specific markers such as Bcl6 and Ret for SMNs shown here and additional cell type specific markers shown in Supplementary Fig. 1.

### Diversity within skeletal motor neurons changes dramatically during maturation

To delineate the maturation trajectory of SMNs from embryo to adulthood, we merged SMNs from all profiled ages and performed a UMAP clustering analysis without integration to preserve differences between ages (**Fig. 2a, 2b**, Methods). This analysis recapitulated several known features of SMNs, underscoring the robustness of our dataset: first, we observe that cells from E15.5 to P21 time points are found in non-overlapping clusters, and that cells from P56 cluster entirely with P21 cells (**Fig. 2a**). Our previous bulk RNA-seq analysis of SMNs showed that SMNs change progressively as they mature from E10.5 to P21, after which time gene expression remains largely stable^5^. Consistent with these previous findings, our current results show that P21 and P56 SMNs have similar gene expression profiles at the single nucleus level, even though these datasets were generated from SMNs collected and processed separately. Second, we find that E15.5 SMNs segregate into dozens of transcriptionally distinct clusters. These clusters have gene expression patterns that correspond to established column/pool identities that have been identified in the embryo^1^ (**Fig 2c, Supplementary Fig. 2b, 2c**) and are discussed in more detail in the next section. Third, we note that the P21/P56 cells are found mainly in three distinct groups. We confirmed that these groups correspond to alpha, gamma, and type3 subtypes based on markers identified in previous single nucleus RNA-seq studies of adult SMNs^19,20^ (**Fig. 2d**).

**Figure 2:**
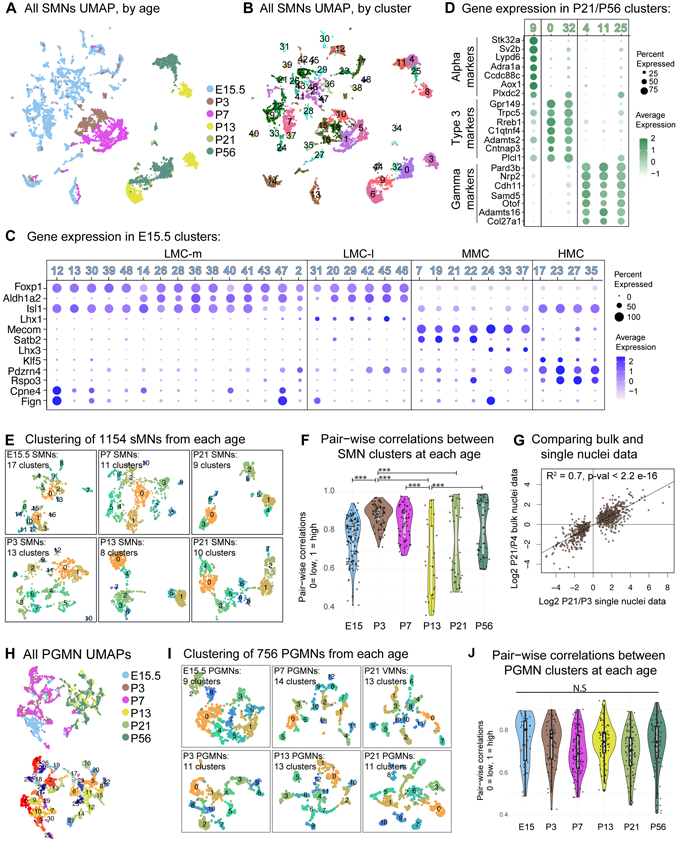
SMN diversity fluctuates during maturation. **A, B)** UMAP representation of SMNs at all profiled ages, colored by age (A) and cluster identities (B). **C)** Gene expression in E15.5 clusters from UMAP in (A) and (B). E15.5 clusters divide into known motor column identities including Lateral Motor Column, which can be subdivided into lateral and medial (LMC-m, LMC-l), Medial Motor Column (MMC), and Hypaxial Motor Column (HMC). **D)** Gene expression in P21/P56 clusters from UMAP in (A) and (B). These clusters express markers of alpha, gamma, and type3 identities in mutually exclusive patterns. **E)** UMAP clustering of 1154 randomly selected SMN nuclei from each age. **F)** Pair-wise Pearson correlations between age-specific clusters from (E). Each dot represents a single pair-wise correlation, with barplot and violin showing the distribution of data. P-value from ANOVA with Tukey’s correction, *** = p-value < 1e-6. **G)** A comparison of differential gene expression from published bulk data (P4 vs. P21) and single nucleus data (P21 vs. P3). Lm model was used for regression. **H)** UMAP representation of PGMNs at all profiled ages colored by age (top) and cluster identities (bottom). **I)** UMAP clustering of 756 randomly selected PGMN nuclei from each age. **J)** Pair-wise Pearson correlations between age-specific clusters from (I). Each dot represents a single pair-wise correlation, with barplot and violin showing the distribution of data. P-value from ANOVA with Tukey’s correction, NS = p-value >=0.5.

Having recapitulated these known features, we next turned to the novel maturation trajectory revealed by this dataset. The snMultiome data showed that the diversity within SMNs fluctuates dramatically over time (**Fig. 2a, 2b**). While E15.5 SMNs are highly diverse and can be segregated into dozens of distinct clusters, these are no longer present one week later at P3. By P3, SMNs collapse into a transcriptionally more homogenous state, but they diversify again in the first two weeks of postnatal life into the three alpha, gamma, and type3 identity clusters found at P13, P21 and P56. To evaluate the molecular diversity among SMNs at each stage more rigorously, we performed a correlation analysis between nuclei at each individual time point. This was done by i) independently clustering 1154 randomly selected SMNs at all six timepoints, which resulted in 8-17 clusters per age (e.g., 17 clusters at E15.5,13 clusters at P3) (**Fig. 2e**), and ii) calculating the pairwise Pearson correlations between clusters within each age based on the expression of top marker genes (**Fig 2f**, Methods). This analysis showed that while E15.5 clusters showed a range of low and high correlations (ranging from 0.41 to 0.94), the clusters at P3 were significantly more correlated to one another (with correlation ranging from 0.72 to 0.97). The correlations between clusters spread out again by P13, reaching a range that is similar to E15.5 (0.36 to 0.96). This analysis therefore reinforces our initial observation of an hourglass shaped maturation trajectory: SMNs are highly diverse at E15.5, their differences coalesce a week later at neonatal stage P3, and they diversify again in the first two weeks of life. Lastly, to ensure that the lack of diversity at P3 is not because of poor data quality, we performed differential gene expression between P3 and P21 SMNs from our current dataset and compared it to differential gene expression between our previously published P4 and P21 bulk RNA-seq data from SMNs^5^. This analysis showed a significant correlation between the two datasets (R^2^ = 0.7, p-val < 2.2e-16), despite technical differences between bulk nuclei and single nucleus RNA-seq (**Fig. 2g**).

To determine if an hourglass trajectory is a shared feature of all spinal motor neurons, we examined the cholinergic PGMNs in our dataset. Recent single nucleus studies have shown that adult PGMNs are highly diverse^19,20^, and we asked if the diversity within these cells also changes over time by performing the same analyses as SMNs: joint UMAP of all ages, and correlation analysis at individual ages (**Fig. 2h-2j**). The joint UMAP of PGMNs from all ages showed that PGMN nuclei divide into two main groups: one that contains cells from E15.5, P3, and P7, and another that contains cells from P13, P21, and P56 (**Fig. 2h)**, suggesting that unlike SMNs which form largely distinct clusters from E15.5 to P21, PGMNs undergo a major transcriptional shift only between P7 and P13. And, we also observe that PGMN diversity does not fluctuate over time: PGMNs of all ages divide into similar numbers of clusters in the joint UMAP analysis (**Fig. 2h**) and they demonstrate a similar range of correlations at each individual timepoint (**Fig. 2i, 2j**).

Overall, these data suggest that SMNs follow a novel and thus far unique developmental trajectory which involves diversification into embryonic subtypes, the loss of embryonic diversity during perinatal life, and the re-diversification into adult subtypes. We next took a closer look at SMN diversity in the embryo and adult, and the transitions between timepoints.

### Extensive column and pool diversity of embryonic SMNs is lost during maturation

Developmental studies in mouse, chick, and human embryos demonstrate that embryonic SMNs divide into spatially restricted categories called columns, which further divide into pools. In the following series of analyses, we confirm that the E15.5 nuclei in our dataset reflect known patterns of embryonic diversity, and then examine whether any aspects of this diversity are maintained to adulthood.

To understand the degree to which E15.5 SMNs reflect known motor column and pool diversity, we performed UMAP clustering on all E15.5 SMNs in our dataset. This led to the identification of 42 clusters (**Supplementary Fig. 2a**) which represent all five described SMN columns. 25 of the clusters express Lateral Motor Column (LMC) marker Foxp1 at high levels and further divide into lateral and medial LMC based on mutually exclusive expression of Lhx1 or Isl1^26^. 10 clusters express the Medial Motor Column (MMC) marker (Mecom), 5 clusters express Hypaxial Motor Column (HMC) markers (Foxp1-low, Isl1+), 1 cluster expresses Phrenic Motor Column (PMC) markers (Pou3f1 +, Chst9+, Klf5 +), and 1 small cluster expresses Spinal Accessory Column (SAC) markers (Runx1+, Phox2b+)^1^ (**Supplementary Fig. 2b, 2c**). In addition to motor column identities, marker expression suggests that individual clusters within larger columns represent distinct motor pools. Among limb innervating LMC neurons, 6 clusters are FoxP1+, Isl1+, but do not express Aldh1a2 (Lm1-Lm6), likely representing digit innervating motor pools. Out of these 6 clusters, Lm1 expresses markers of intrinsic digit innervating pools: Fign+, Cpne+, Pou3f1+; two additional clusters (Lm3, Lm6) express markers associated with extrinsic flexor innervating pools: Pou3f1+, Fign-, Cpne-^36^. We can also identify individual clusters that express previously described pool specific TFs Etv1 (or Er81 in L, H3), and Etv4 (or Pea3, expressed in Lm2, Lm7, Ll3)^1,37^. Lastly, we observe that MMC neurons can be clearly divided into a Satb2+ population and a Nr2f2+ population, as recently described in mouse E13.5 and human embryos^21^ (**Supplementary Fig. 2c**). The embryonic neurons in our dataset therefore reflect the rich diversity that has previously been described by genetic and single cell studies. We add to these studies by identifying novel column/pool specific genes (**Supplementary Table 1**). In addition, because our data contains ATAC-seq, we have identified column/pool specific accessible chromatin regions that will provide novel insights for future studies of embryonic SMNs (**Supplementary Table 2**).

The diversification of embryonic SMNs into columns and pools ensures that SMN axons reach their specific muscle targets. Once this SMN to muscle circuitry is formed in the embryo, it is maintained throughout adulthood. We therefore wondered if gene expression signatures of column and pool diversity may also persist in adults, even though the clustering of mature neurons seems to be dominated by alpha vs. gamma. vs. type3 identities. To investigate this, we first examined the average expression of column and pool specific gene modules – consisting of the top 100 marker genes for each column (LMC, MMC, HMC) or top 20 marker genes for pools (individual clusters from Supplementary Fig 2a) – at all profiled ages. This analysis showed that column and pool gene modules are not maintained as SMNs undergo postnatal maturation. Expression of column modules is clearly seen in specific embryonic clusters at E15.5, but by P13 these signatures are largely downregulated (**Fig. 3a-3c**). The proportion of SMNs that express pool modules also decreases to almost zero by P13 (**Fig. 3d**).

**Figure 3.**
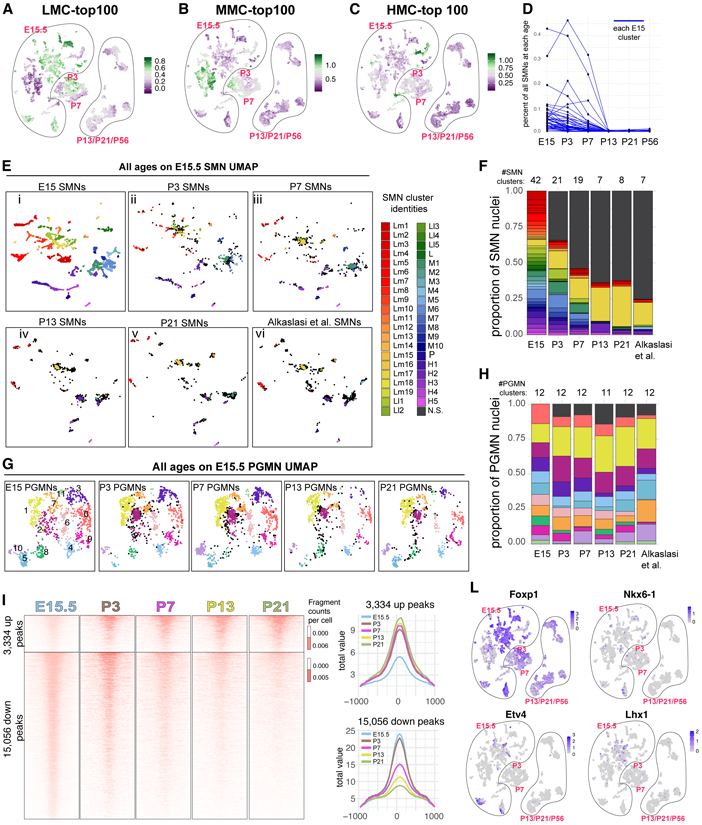
Embryonic SMN diversity is lost during maturation. **A, B, C)** Average expression of LMC, MMC, and HMC gene modules (top 100 genes) in all SMNs during maturation. More green denotes high expression, more purple denotes low expression. Gene lists are in Supplementary Table 1 for a-d. **D)** Proportion of SMNs at each age that express gene modules (top 20 genes) of each of the E15.5 clusters from Supplementary Fig. 2a. Each blue line shows expression of one cluster. **E)** SMNs from all ages, and from a previously published dataset (Alkaslasi et al., 2021) are plotted on the E15.5 UMAP using label transfer analysis in Seurat. Colored dots are SMNs that show a significant match to E15.5 clusters. Dark gray dots are nuclei that do not show a significant match to E15.5 clusters. **F)** Proportion of SMNs from each age that do or do not show a significant match to E15.5 clusters based the label transfer analysis in (E). Each color represents the matching cluster from (E). **G)** Label transfer analysis on PGMN nuclei. PGMN nuclei from all ages are plotted on the E15.5 PGMN UMAP. Colored dots are PGMNs that show a significant match to E15.5 clusters while dark gray dots are PGMNs that do not. **H)** Proportion of PGMNs from each age that do or do not show a significant match to a specific E15.5 clusters based the label transfer analysis in (G) with colors representing the matching cluster. **I)** Open chromatin regions that show cluster specific accessibility at E15.5. On the left is a heatmap, and on the right line plots, demonstrating changes in ATAC-seq reads at these peaks over time. Up peaks show increased accessibility between E15.5 and P21 (E15.5 < P21) while down peaks show a decrease in accessibility between E15.5 and P21 (E15.5 > P21). **J)** Expression of Foxp1, Etv4, Nkx6.1, and Lhx1 during maturation. These TFs are known to regulate column and pool identity specification in the embryo.

As a complementary approach, we used label transfer in Seurat to determine if SMNs from P3, P7, P13, and P21 retain transcriptional differences that define pool and column clusters found in the E15.5 UMAP^38^. In this analysis, SMNs from different ages are independently plotted on the E15.5 UMAP and assigned the E15.5 identities that they are closest to. We found that both the proportion of SMNs with a significant match to E15.5 clusters, and the number of E15.5 clusters that they are matched to, decreases over time. At P3, 66% of SMNs show a significant match to E15.5 clusters, and these SMNs subdivide into 21 of the 42 E15.5 clusters. By P21, this dropped to 37% of SMNs which mapped to only 8 clusters (**Fig. 3e, 3f**). To further validate these findings, we performed the same label transfer analysis on a published snRNA-seq dataset from adult Chat-Cre>SUN1-GFP SMNs (Alkaslasi et al. 2021^20^, Methods). When the Alkaslasi et al. adult SMNs are plotted on our E15.5 UMAP space, we see very similar results – 25% of adult SMNs show a significant match to E15.5 clusters, and these nuclei segregate into 7 of the 42 E15.5 clusters (**Fig. 3e(vi), 3f**). Notably, mature SMNs that continue to show significant matches to the 7-8 E15.5 clusters seem to congregate at the center of LMC, MMC or HMC clusters, suggesting that while pool identities are likely lost over time, some aspects of column identities may be retained.

To determine if the loss of embryonic diversity is specific to SMNs, we performed a similar label transfer analysis on the PGMNs in our dataset. We observe that >90% of P21 PGMNs retain a significant match to E15.5 clusters (compared to 37% of SMNs), and these cells divide into 12 of 12 identified E15.5 clusters. 92% of adult PGMNs from Alkaslasi et al. also show a significant match to E15.5 PGMNs in our dataset and divide into all 12 clusters (**Fig. 3g, 3h**). These results demonstrate that the loss of embryonic diversity is a specific feature of the SMN maturation trajectory and is not observed in PGMNs.

Transient molecular events, such as signaling gradients during early development or neural activity, can be “remembered” in cells through epigenetic changes, such as DNA methylation or histone modification, and influence gene expression or function at later times^39,40^. We asked whether maturing SMNs retain a similar memory of embryonic pool diversity in their open chromatin signature even though cluster specific gene expression patterns are transient. To address this, we identified 19,048 ATAC-seq peaks that show cluster specific accessibility patterns at E15.5 and asked if these peaks maintain accessibility over the course of maturation. We found that 79% of E15.5 cluster specific peaks lose accessibility over time, 17.5% gain accessibility over time and the remaining 3.5% remain unchanged (**Fig. 3i**, **Supplementary Fig. 3a, 3b**). We next examined if the 17.5% of peaks that gain accessibility over time are maintained in discrete populations of cells where they may serve as an epigenetic memory, but found that these peaks do not remain enriched in specific cells. Rather, they show diffuse accessibility in a larger proportion of cells over time (**Supplementary Fig. 3c)**. These data therefore demonstrate that maturing SMNs do not maintain the chromatin accessibility signatures that differentiate SMN pools and columns.

Finally, we examined the expression pattern of column and pool specific TFs over the course of maturation. Cell intrinsic TFs regulate both gene expression and chromatin accessibility and have been shown to be important for establishing column and pool identities. We identified 74 TFs that show cluster enriched expression at E15.5 and found that 60% of these are downregulated over time (**Supplementary Fig. 3d**), 11% are upregulated (**Supplementary Fig. 3e**), and the remaining 29% do not change significantly (**Supplementary Fig. 3f**). TFs that are downregulated include factors that have previously been identified to have roles in pool and column specification such as Foxp1, Etv4 (Pea3), Nkx6-1, Isl1, Lhx3, Lhx1, Pou3f1 (Scip), and Hox TFs^1^ (**Fig. 3j**, **Supplementary Fig. 3d**). TFs that are upregulated, such as Thrb and Mitf (**Supplementary Fig. 3e, 3g**) become less specific over time and are expressed in a higher proportion of SMNs, suggesting that they may no longer control pool/column-specific gene expression patterns and may serve new functions. Overall, these analyses show that a vast majority of TFs do not maintain cell type specific expression and support the conclusion that embryonic diversity is largely not maintained as SMNs mature: at the levels of gene expression, chromatin accessibility, and transcriptional regulation.

### Adult alpha, gamma, and type3 identities are built sequentially during maturation

Having established that embryonic diversity collapses during maturation, we next focused on the emergence of adult diversity. To determine when SMNs begin to show adult subtype identities, we plotted the average expression of adult subtype gene modules – consisting of top 50 markers of mature alpha, gamma, and type3 identities – at all profiled ages. This analysis revealed that alpha and gamma modules show very low expression in the embryo, but become more highly expressed and more selective during postnatal life (**Fig. 4a, 4b**). The Type3 module, on the other hand, is not expressed at all in the embryo and becomes activated between P3 and P7 in a selective manner (**Fig. 4c**). To understand the timing of subtype development in more detail, we used label transfer analysis to plot SMNs from E15.5, P3, P7, and P13 on the P21 UMAP, where SMNs are clearly divided into three alpha, gamma, and type3 clusters (clusters P21-c2, P21-c0, and P21-c1 respectively) (**Fig. 4d-4e**). This analysis shows that 58% of E15.5 and 88% of P3 SMNs show a significant match to P21 clusters, however these cells only segregate into alpha and gamma identities. By P7, 97% of SMNs show a significant match to P21 clusters and these cells segregate into alpha, gamma, and type3 identities (**Fig. 4d, 4f**). 99% of adult SMN nuclei from Alkaslasi et al., also map on to alpha, gamma, and type3 clusters in our P21 UMAP, demonstrating the congruence between our datasets (**Fig. 4d, 4f)**. Thus far, both the expression of gene modules and label transfer analyses suggest that alpha and gamma identities emerge around E15.5-P3, but type3 identity emerges later between P3 and P7.

**Figure 4.**
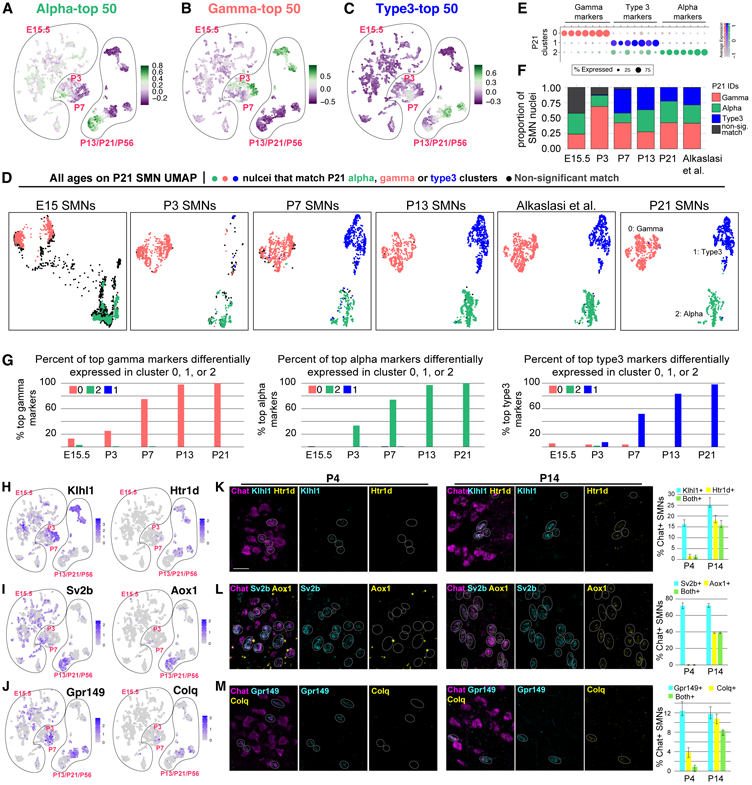
Adult SMN diversity is acquired during maturation. **A, B, C)** Expression of alpha, gamma, and type3 gene modules (50 genes) in all SMNs during maturation. Gene lists are in Supplementary Table 3. **D)** SMNs from all ages and from Alkaslasi et al. are plotted on the P21 UMAP using label transfer analysis in Seurat. SMNs that show significant match to P21 cluster 0 (red), 1(blue) or 2 (green) are colored; SMNs that do not show a significant match are dark gray. **E)** Clusters 0, 1, and 2 from P21 correspond to gamma, type3, and alpha identities respectively. **F)** Quantification of proportions of SMNs from each age and from Alakaslasi et al., that do or do not show a significant match to P21 alpha, gamma, and type3 clusters based on label transfer analysis. Colors match cluster colors in (D). **G)** Percent of top gamma, alpha, and type3 markers that are differentially expressed in clusters 0, 8, and 2 at each age. **H, I, J)** Expression of early and late markers of gamma (H), alpha (I), and type3 (J) identities in the multiome data from all SMNs. **K, L, M**) RNA FISH validation of early and late marker expression in P4 and P14 spinal cords. Images show ventral lateral region of the spinal cord where all Chat+ cells are SMNs. Each circle outlines a single cell. Colors of FISH expression matches gene names. Quantification of marker expression as percent of Chat+ SMNs is shown on right.

We next asked if all genes that show selective expression in mature alpha, gamma, and type3 SMNs are activated at the same time. For example, do SMNs that segregate into alpha and gamma clusters at E15.5 and P3 already express all alpha and gamma specific genes, or do they acquire their gene expression pattern over the course of maturation. To answer this, we examined the expression of subtype marker genes in E15.5, P3, P7, and P13 SMNs that are predicted to belong to alpha (P21-c2), gamma (P21-c0), or type3 (P21-c1) clusters (**Fig. 4g)**. We found that not all top alpha, gamma, and type3 genes are activated at once. Instead, in all three subtypes, gene expression programs are temporally regulated. Some subtype specific genes such as Klhl1, Sv2b, and Gpr149 are activated early, some like Htr1d, Colq, and Aox1 are activated later (**Fig. 4g, Supplementary Figure 4a-4c**). Additionally, we identified genes like Cdh11, C1qtnf4, and Ctcn5 that are broadly expressed early on, but become restricted to a specific subtype in mature animals (**Supplementary Figure 4d)**. Notably, we found that subtype specific gene expression emerges in a sequential manner: gamma genes start to show selective expression (Log2FC > 1) as early as E15.5, followed by alpha at P3, and then type3 by P7. Gamma genes that are already differentially expressed at E15.5 include Creb5, Htr1f, Klhl1, Plekhg1. Some alpha genes, like Chodl and Pld5, start to show differential expression at E15.5, but this does not reach the threshold of Log2FC > 1. Overall, this data demonstrates that mature gene expression patterns are acquired over time through both activation and repression of genes and that gamma, alpha, and type3 identities are specified in that order between E15.5 and P7.

To validate the sequential activation of subtype genes, we performed HCR RNA FISH at P4 and P14 for early and late activated alpha, gamma, and type3 genes. We used Klhl1 and Htr1d as early and late gamma markers, Sv2b and Aox1 as early and late alpha markers, and Gpr149 and Colq as early and late type3 markers respectively (**Fig. 4h-4j**). As predicted by our single nucleus data, we found that Klhl1, Sv2b, and Gpr149 are already expressed in Chat+ SMNs at P4 and are not significantly upregulated between P4 and P14 (**Fig. 4k-4m**, left P4 panels). In contrast, Htr1d, Aox1, and Gpr149 are significantly upregulated between P4 and P14 (**Fig. 4n-4m**, right P14 panels and bar plot). We note that the proportions of alpha (∼70%), gamma (∼18%), and type3 (∼12%) observed by RNA FISH are similar to previous in vivo reports^41^, but do not reflect the proportions observed in our single nucleus multiome data (**Fig. 4f**), likely due to technical hurdles in isolating alpha nuclei which are larger and potentially more fragile. Overall, our single nucleus sequencing data and RNA FISH validation suggests that distinct alpha, gamma, and type3 subtypes become evident by neonatal life, and that these subtypes continue to acquire subtype specific gene expression programs till they become fully mature in the third week of life.

### Alpha, gamma, and type3 identities are regulated by temporally activated chromatin regions and TFs

We next asked how adult subtype gene expression patterns are acquired over a 3-4 week time period. Regulatory regions that drive adult-subtype gene expression may already be primed and accessible in the embryo, or they may become accessible over time, through the action of temporally activated TFs. To distinguish between these possibilities, we identified alpha-, gamma-, and type3-specific peaks at P21/P56 and then looked at ATAC-seq reads at these regions over time. Differential ATAC analysis at P21/P56 led to the identification of ∼12,000 gamma enriched peaks, ∼20,000 alpha enriched peaks, and ∼10,000 type3 enriched peaks, which we filtered to 4071 gamma peaks, 2987 alpha peaks, and 2301 type3 peaks based on stricter criteria (logFC > 2, and adjusted p-value < 0.001) (**Fig 5a**). By plotting the ATAC-seq reads at these peaks over the course of maturation, we found that subtype-specific peaks were largely inaccessible at E15.5, but become accessible during postnatal life (**Fig. 5b**). Gamma-specific open chromatin regions become accessible earliest, reaching a P21-like state by P3, whereas alpha and type3 open chromatin regions become accessible more gradually, in agreement with earlier acquisition of subtype specific gene expression patterns in gammas compared to alpha and type3 (**Fig. 5b**, lineplots on right, **Fig. 4g**).

**Figure 5:**
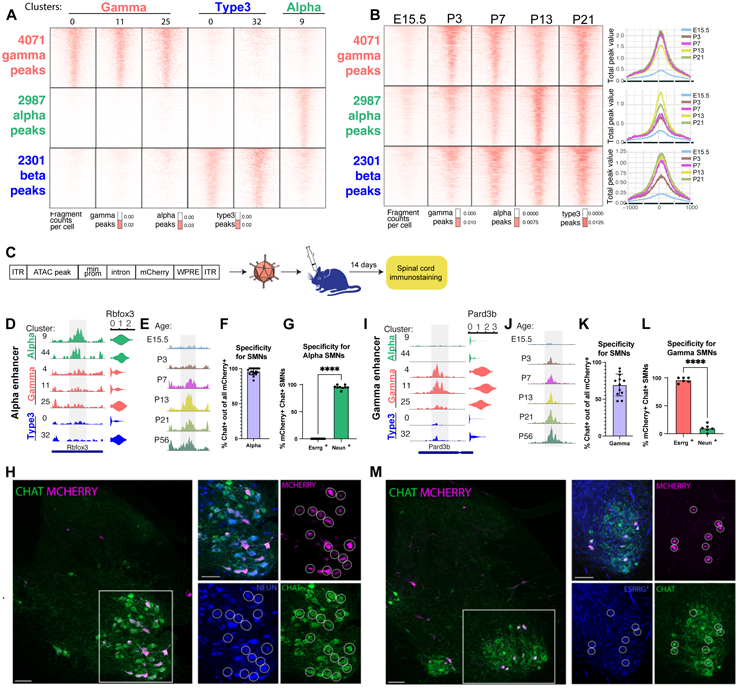
Subtype-specific chromatin regions are acquired over time and drive subtype-specific gene expression. **A, B)** Heatmaps of snATAC-seq reads at alpha, gamma, and type3 specific open chromatin regions in mature P21/P56 SMN clusters from Figure 2a, 2b (A), and in all SMNs of different ages (B). In (B), summary lineplots are presented on the right. **C)** Schematic of enhancer-AAV experiment. **D, E)** Alpha-enriched open chromatin regions shown in mature clusters (D) and at all ages (E). **F)** Percent of AAV-AphaEnhancer-mCherry expressing cells that overlap with CHAT+ SMNs in the ventral horn of the spinal cord. **G)** Percent of AAV-AlphaEnhancer-mCherry expressing cells that overlap with alpha marker NEUN and gamma marker ESRRG. **H)** Images demonstrating co-expression of AAV-AlphaEnhancer-mCherry with CHAT and NEUN. Each circle is a single mCherry+ cell. **I, J)** Gamma-enriched open chromatin regions shown in mature clusters (I) and at all ages (J). **K)** Percent of AAV-GammaEnhancer-mCherry expressing cells that overlap with CHAT+ SMNs in the ventral horn of the spinal cord. **L)** Percent of AAV-GammaEnhancer-mCherry expressing cells that overlap with alpha marker NEUN and gamma marker ESRRG. **M)** Images demonstrating co-expression of AAV-GammaEnhancer-mCherry with CHAT and ESRRG. Each circle is a single mCherry+ cell. *ESRRG is an anti-Mouse primary antibody and shows background labeling.

We next functionally tested the ability of temporally activated open chromatin regions to drive subtype specific gene expression using enhancer reporter assays. To do this, we identified alpha and gamma-specific open chromatin regions that are linked to alpha and gamma specific genes respectively (**Fig. 5d, 5i**), and that become accessible after E15.5 (**Fig. 5e, 5j)**. We generated AAV-enhancer-mCherry constructs with these open chromatin regions, produced AAVs using published protocols, injected mice retro-orbitally at P21-P35, and checked expression of mCherry in the spinal cord 14 days later (**Fig. 5c**). We tested three candidate enhancers that show alpha or gamma specific accessibility and followed up with one alpha and one gamma enhancer that showed consistent expression in Chat+ SMNs (**Fig. 5d, 5f, 5i, 5k**). To test the alpha vs. gamma specificity of these enhancers, we performed immunostaining for NEUN/RBFOX3 and ESRRG, markers that have previously been validated as alpha and gamma enriched respectively^22^. AAVs containing the alpha specific enhancer showed mCherry expression in NEUN+ cells, but not in ESRRG+ cells (**Fig. 6h**). Conversely, AAVs containing the gamma enhancer expression in ESRRG+ SMNs but not NEUN+ SMNs (**Fig. 6m**). This data demonstrates that temporally activated, subtype-specific open chromatin regions do regulate subtype specific gene expression.

**Figure 6:**
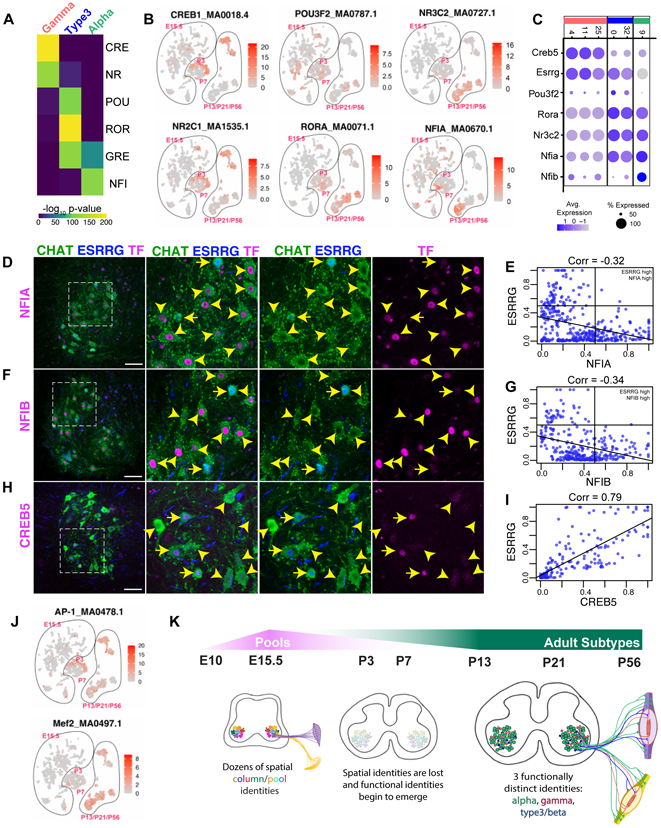
Identification of candidate TFs that regulate adult subtype specification. **A)** Heatmap of TF motif enrichment in subtype-specific open chromatin regions from Figure 5A. **B)** Enrichment of subtype-specific TF motifs in ATAC-seq peaks of all SMNs during the course of maturation. Jaspar motif names are shown in addition of motif names. **C)** snRNA-seq expression of specific TFs that bind to motifs shown in (A) and (B). **D, F, H)** Immunostaining of candidate TFs with ESRRG, a TF previously shown to have gamma-enriched expression. Arrow heads point to ESRRG-SMNs, while arrows with stems point to ESRRG+ SMNs. ESRRG is an anti-Mouse primary antibody and shows background labeling. **E, G, I)** Quantifications of immunostainings to the left. Each dot represents one cell, with relative fluorescence of each TF on x-axis and relative fluorescence on ESRRG on y-axis. Pearson correlation of pairs of TFs analyzed are on top. **J)** Enrichment of activity-dependent TF motifs in ATAC-seq peaks of all SMNs during the course of maturation. Jaspar motif names are shown in addition of motif names. **K)** Model demonstrating the changes in SMN diversity over the course of maturation.

To identify potential transcriptional regulators of subtype identity, we performed motif enrichment analyses in subtype-specific peaks. This led to the identification of unique motif combination in each subtype: CRE (Creb response elements) and NR (nuclear receptor) motifs were specifically enriched in gamma-specific peaks, ROR (RAR-related orphan receptor) and POU motifs were specifically enriched in type3 SMNs, GRE (glucocorticoid response element) motifs were enriched in both type3 and alphas, and NFI motifs were enriched specifically in alpha SMNs (**Fig. 6a**). When we looked at the prevalence of these motifs in ATAC-seq data from all SMNs (as opposed to just in subtype-specific peaks), we also observed subtype specific enrichment in mature P13/P21/P56 clusters. Interestingly whereas CRE and NFI show specific enrichment by P3, ROR becomes specifically enriched by P7 and GRE by P13 (**Fig. 6b)**. This suggests that activation of regulatory TFs may also follow a temporal pattern, similar to subtype-specific gene expression and chromatin accessibility.

Our previous bulk ATAC-sequencing of SMNs had shown enrichment of activity-dependent AP-1 and Mef2 motifs in mature SMNs. Although these motifs were not highly enriched in subtype-specific peaks analyzed above, they did show maturation dependent enrichment in all SMN subtypes. Mef2 motifs are highly enriched in all subtypes at P13/P21/P56, while the AP-1 motif showed an interesting temporal pattern-gamma SMNs are enriched for AP1 motifs already at P3, alpha SMNs at P13, and type3 SMNs at P21/P56 (**Fig. 6j**). This is consistent with the temporal sequence of maturation we have observed so far: gamma SMNs first, followed by alpha, and then type3.

TF DNA-binding domains are often shared by a whole family of TFs, we therefore looked for family members that showed subtype-enriched expression and were able to identify TFs that matched each of the motifs: Creb5 (CRE-binding TF) and Esrrg (NR-binding TF) are differentially highly expressed in gamma SMNs. Nr3c2 (GRE-binding TF) and Rora (ROR-binding TF) are expressed in both alpha and type3 SMNs, Pou3f2 (binds POU motifs) is differentially expressed only in type3 SMNs, and Nfia + Nfib (NFI-binding TF) are differentially more highly expressed in alpha SMNs (**Fig. 6c**). We have previously shown that Nr3c2 expression at the protein level is activated in SMNs between P4 and P21^5^, which is consistent with GRE motifs showing enrichment between P7 and P13 and further suggests Nr3c2 is a possible regulator. Here we validated the subtype specificity of three additional TFs (Nfia, Nfib, and Creb5) by co-staining with previously validated gamma marker ESRRG. As predicted by gene expression and motif enrichment, we found that both NFIA and NFIB are more highly expressed in ESRRG- neurons than in ESRRG+ neurons. Conversely, CREB5, which has previously not been studied in the context of SMNs, shows higher expression in ESRRG+ gamma SMNs (**Fig. 6d – 6i**). These TFs are therefore strong candidates for regulating subtype specific gene expression in adult SMN subtypes. Altogether, our work supports a model in which subtype-specific transcriptional regulators may work with shared activity-dependent factors to control the step-by-step acquisition of adult SMN subtypes during maturation.

## Discussion

Our results show that diversity within SMNs fluctuates dramatically as they transition from embryonic to neonatal to mature stages (**Figure 6k**). In the embryo, when SMN circuits to specific muscles are being wired, SMNs divide into dozens of spatial pool and column identities. After SMN axons have reached their muscle target, the gene expression pattern and chromatin accessibility that defines these identities are lost, even though the SMN to muscle circuits are maintained throughout life. The loss of embryonic transcriptional diversity is accompanied by a re-diversification of SMNs into the three main transcriptionally- and functionally-distinct alpha, gamma, and type3/beta subtypes which are then maintained into adulthood. These subtypes innervate different types of fibers within each muscle and have distinct electrophysiological properties that ensure proper muscle contraction and stretch. Our work demonstrates that transcriptional diversity within a neuron type changes over time to reflect the functional needs of the cells. These trajectories can therefore take different shapes depending on the sequence of events that each neuron undergoes to reach its functional, adult state.

### A novel maturation trajectory

A handful of studies have now used single cell sequencing to map the developmental trajectories of mammalian neurons from embryo to adulthood. The most common maturation trajectory to emerge from these studies is one in which diversity within a neuronal population increases over time. Single cell RNA-seq of mouse dorsal root ganglia sensory neurons, human and mouse cortical interneurons, and human cortical excitatory neurons all show that diversity within these neuron types increases during maturation as subpopulations acquire distinct transcriptional and functional features over time^6,9,14,15,42,43^. In the preoptic area of the mouse hypothalamus, a different maturation trajectory is observed where diversification of excitatory and inhibitory neuronal subtypes is apparent already in the embryo, soon after neurogenesis, and this diversity is maintained through adulthood even though these neurons continue to undergo gene expression changes^7^. This pattern is similar to one we observe in the PGMNs in our current study. In maturing SMNs, we have discovered a third and novel maturation trajectory with fluctuating levels of diversity-embryonic SMNs diversify into dozens of spatial identities, this diversity is lost by perinatal stages, and then functional diversity is gained in the first three weeks of life which is maintained during adulthood.

Embryonic SMN diversity required for correct muscle targeting is acquired between ∼E10.5-E15.5 in mouse embryos and then lost during perinatal stages^21,44,45^. This phenomenon of increased diversity specifically during the period of axon targeting has also been noted in drosophila olfactory and optic lobe neurons^46,47^. For example, in the drosophila optic lobe the direction sensing T4 and T5 neurons subdivide into four subtypes between pupal stages 15 and 50, when these neurons are undergoing axonogenesis and synaptogenesis. Once axons have innervated their targets, T4/T5 neurons lose their subtype identities and become homogenous. Unlike mouse SMNs, which re-diversify into adult subtypes after losing embryonic diversity, T4/T5 neurons remain in this homogenous state throughout adulthood. This may be because once T4/T5 neurons have correctly reached their respective targets, they perform very similar functions throughout life. On the other hand, once SMNs have reached their correct muscle targets, they undergo a second round of diversification to acquire specific electrophysiological properties that allow them to correctly control extrafusal muscle contractions, intrafusal fiber stretch, or both. This phenomenon of transient increases in diversity during early stages of development may also explain why the multimodal single cell profiling by the Brain Initiative Cell Census Network revealed more fine-grained morphological diversity than transcriptional diversity among adult projection neurons^48^. The gene expression programs required to generate the fine-grained morphological diversity may have been activated transiently when these projection neurons were forming circuits, but downregulated by adulthood after circuits are established. These studies support a model in which the developmental maturation trajectory of a neuron is driven by its changing functional needs. Thus, neuronal identity and diversity can look very different at different time points, and adult neurons are best understood as functional outputs of transcriptional events across their maturation timescale.

### Timing of adult SMN subtype diversification

We found that alpha, gamma, and type3 subtypes acquire their adult-like gene expression patterns over the course of ∼3 weeks from perinatal to juvenile stages. Behavioral studies have shown that the acquisition of adult like movement also occurs by the third week of postnatal life^17^, strongly suggesting that the generation of functional identities that ensure proper muscle contraction leads to maturation of motor behaviors. This process of transcriptional diversification in postnatal life is much slower compared to the initial diversification of SMNs into dozens of column/pool identities which occurs over ∼3-5 days. Interestingly, postnatal subtype maturation in this longer timeframe coincides with the maturation of upstream circuits and downstream muscle targets: i) Descending projections form the corticospinal tract reach the lumbar spinal cord around P9-P10 with full innervation occurring around P14^49^. ii) Serotonergic signaling from the brainstem projections to the spinal cord become adult-like at P21^50^. iii) Studies in the mouse EDL muscle have shown that the number of nuclei per myofiber increases 5x between P3 and P21, stabilizing after this point, even though the myofibers themselves continue to grow larger till P56 as animals get bigger^51^. iv) Neuromuscular junctions (SMN synapses with muscles) reach maturity around P21 – at the muscle post-synapse, acetylcholine receptor cluster transition from a plaque-like uniform distribution to a pretzel-like branched structure along with structural and gene expression changes^52^. v) In the SMN presynapse, pruning and refining of axon terminals takes place between P0-P21 as each motor axon makes ∼10x less connections with muscle fibers in mature animals than during perinatal life^53^. These parallel changes in upstream motor circuits and downstream muscle raise interesting questions about interdependencies in SMN subtype maturation. In addition to TFs identified in our multiome analysis, do signals from descending circuits and from maturing muscles play a role in refining adult SMN subtype identity? And conversely, do maturing SMNs provide instructive clues for muscle fiber maturation? In future work, datasets generated here in conjunction with published single nucleus profiling of muscles may help untangle the complex interplay of signals required for correct maturation of both SMNs and muscles.

### Activation of subtype identity in a step-by-step manner

A common mode of cell type specific gene regulation in neurons is through the action of terminal selector TFs^54^. These TFs are generally expressed in neurons throughout their lifetimes and are required to both activate and maintain neuron-type specific gene expression programs. However, our work suggests that sMNs follow a more step-by-step mode of gene regulation which tracks with the changes in diversity that occur over time. The initial specification of SMN identity is controlled by the LIM-homeodomain TFs, Isl1 and Lhx3, which are known to be downregulated or turned off in many SMNs within 1-2 days as cells activate columns/pool-specific TFs^5,29^. Our single cell data now show that a majority of TFs that show column/pool specific expression at E15.5 are also downregulated by P3, including known column/pool TFs such as Etv1, Etv4, Pou3f1^55,56^. And, we have identified a third set of TFs that are enriched in specific adult SMN subtypes and maintained throughout adulthood. These include potential subtype-specific regulators of adult identities: Creb5, Nfia, Nfib, Nr3c2, Rora, Pou3f2, and shared activity dependent TFs. Intriguingly, some of these TFs are activated early (such as the gamma TF Creb5) while others are activated later (such as steroid hormone dependent TF Nr3c2 and activity dependent TFs AP-1 and Mef2), suggesting a temporality to their function

To date, our understanding of the developmental mechanisms that drive adult SMN subtype specification has remained cursory. Gamma-specific functional properties have been shown to be regulated by TFs Err2/Err3^22^. Loss of TFs Err2/Err3 leads to changes in biophysical and firing properties of gamma neurons, however gamma identity as well as gamma innervation of muscle spindle remains intact in these mutants^22^. In alpha neurons, a delta like ligand Dlk1, has been shown to specify fast over slow physiology^57^. However, in Dlk1 mutants, overall alpha identity is preserved. In future work, it will be important to functionally test the role of subtype specific TFs identified here and determine if they control broader aspects of alpha, gamma, and type3 identity. It would also be interesting to determine how these factors work with activity-dependent mechanisms and perhaps other signals form muscles to activate adult subtype genes over the course of ∼3 weeks. Understanding how these intrinsic and extrinsic mechanisms coordinate maturation in SMNs may also help to elucidate similar mechanisms in other neuron types throughout the nervous system.

Overall, this work bridges a major gap in our understanding of SMNs by elucidating how they transition from dozens of subtypes in the embryo to three functionally distinct and disease relevant subtypes in the adults. The datasets and tools developed in this study will allow continued functional dissection of SMN development, function, and disease. More broadly, this work demonstrates that neurons can undergo dramatic changes in transcriptional identity as their functional needs change from embryo to adulthood.

## Methods

### Animal handling

#### Single nuclei experiments

All procedures involving mice for these experiments were approved by the Columbia University Medical Center Institutional Animal Care. These experiments were carried on mouse lines that have been previously characterized: SUN1-sfGFP-Myc ^34^(JAX stock#021039) and ChAT-IRES-Cre::SV40pA::Δneo (Chat-Cre)^35^ (JAX stock#031661). For purification of nuclei at all other ages, homozygous SUN1-2xsfGFP-6xMYC mice were crossed to homozygous ChAT-IRES-Cre::SV40pA::Δneo mice and transheterozygous progeny were sacrificed for experiments. For multiome-seq experiment at each age, spinal cord tissue was combined from 6-10 animals. Both males and females were included in each experiment.

#### RNA FISH, Immunostaining and enhancer-AAV experiments

All procedures involving mice for these experiments were approved by the Rutgers University Medical School’s Institutional Animal Care and Use Committee. These experiments utilized WT C57BL/6J (JAX stock#000664) mice. For RNA FISH: 4 animals (mix of both sexes) were used for each probe combination, at least 4 sections from brachial or lumbar spinal cords of each animal were scored. For immunostaining: 3-4 animals (mix of both sexes), and at least 3 lumbar section per animal were imaged and scored for each antibody combination. For AAV experiments: 3 mice (mix of both sexes) and 3 lumbar sections per mice were imaged and scored.

### Single Nuc Seq procedure and analysis

Brachial and lumbar regions of the spinal cord were dissected from 6-10 Chat-Cre> SUN1-sfGFP-Myc mice at each experimental age. Nuclei were isolated and sorted following the exact protocol detailed in Patel et al.^5^ All nuclei collected (∼10,000 per mouse) were taken to the Genome Center at Columbia University for single nucleus multiome analysis using the 10X multiome kit protocol with a 10x Chromium. Each age was processed independently on different days and 10,000 nuclei were targeted for each age. Data was pre- processed using CellRanger by the Genomics Core, and imported for analysis in Seurat. High-quality cells were first filtered using the following parameters: nCount_ATAC<7e4 & nCount_ATAC>5e3 & nCount_RNA<25000 & nCount_RNA>5000. These parameters were determined based on the counts distribution of all samples. We also ensured that UMAP analysis did not lead to clusters made solely of low quality cells. Processing of single nucleus data was performed using the following packages: Seurat, Signac, chromVAR, JASPAR2020, TFBSTools, BSgenome.Mmusculus.UCSC.mm10, EnsDb.Mmusculus.v79.

#### Clustering of all cells at each age

At each age, nuclei that made it through the QC cutoff were clustered using both RNA-seq and ATAC-seq data with FindMultimodalNeighbors with Seurat version 5.0.2. wsnn resolution is 0.8 for all.

1. First RNA-seq was processed using: age.obj <- SCTransform(age.obj, verbose = T) %>% RunPCA() %>% RunUMAP(dims = 1:50, reduction.name = ‘umap.rna’, reduction.key = ‘rnaUMAP_’, return.model = T)
2. Then, ATAC-seq was processed using: age.obj <- RunTFIDF(age.obj) %>% FindTopFeatures(min.cutoff = ‘q0’) %>% RunSVD() %>% RunUMAP(eduction = ‘lsi’, dims = 2:50, reduction.name = "umap.atac", reduction.key = "atacUMAP_", return.model = T)
3. Then multimodal clusters were identified using: age.obj <- FindMultiModalNeighbors(age.obj, reduction.list = list("pca", "lsi"), dims.list = list(1:50, 2:50)) age.obj <- RunUMAP(age.obj, nn.name = "weighted.nn", reduction.name = "wnn.umap", reduction.key = "wnnUMAP_", return.model = T) age.obj <- FindClusters(age.obj, graph.name = "wsnn", algorithm = 3, verbose = FALSE)

Figure 1D, top panel shows DimPlot visualizations of reduction = umap.rna. Clusters are colored based on cell identities determined as follows:

First, a combination of FeaturePlot and VlnPlot was used to determine which clusters express Chat, Slc17a6, and Gad1. Then, Chat+ clusters were given SMN identity based on expression of Slc5a7, Bcl6, Ret, Gfra1, and Tns1. Chat+ clusters were given PGMN identity based on expression of Slc5a7, Fbn2, and Zbn2. Chat+ clusters were given PItx2-IN identity if they expressed Slc5a7 and Pitx2.

#### Joint clustering of SMNs/PGMNs from all ages

For joint clustering of SMNs from all ages, clusters assigned SMN identity at each age were subset, and merged without integration as follows:

mn.allages <- JoinLayers(mn.allages, assay = "RNA")

mn.allages <- SCTransform(mn.allages, verbose = T) %>% RunPCA()

mn.allages <- RunUMAP(mn.allages, dims = 1:50, reduction.name = ‘umap.rna’, reduction.key = ‘rnaUMAP_’, seed.use = 18) %>% FindNeighbors(reduction = "pca", dims = 1:50, verbose = T) %>% FindClusters(resolution = 0.8, verbose =T).

The same method was used to perform joint clustering of PGMNs from all ages.

#### Correlation analysis at each age

The same number of SMNs (1157) or PGMNs (756) from each age were randomly sampled. These cells were clustered with the following at each age:

SCTransform(subset.ageobj, verbose = T) %>% RunPCA(return.model=TRUE) %>% RunUMAP(dims = 1:50, reduction.name = ‘umap.rna’, reduction.key = ‘rnaUMAP_’) %>% FindNeighbors(reduction = "pca", dims = 1:50, verbose = T) %>% FindClusters(resolution = 0.8, verbose =T).

Then, at each age, the top 20 marker genes of each cluster and their average expression in all clusters was determined. Pearson correlations were calculated between all cluster pairs based on these average gene expression patterns. For determination of p-values, an Anova test was performed followed by pair-wise comparisons with Bonferroni corrections.

#### Comparison of bulk vs. single nucleus sequencing

Differential gene expression determined by EdgeR on bulk P21 vs. P4 data from Patel et al., 2022 was compared to Differential gene expression between P21 and P3 determined by FindMarkers in merged object of all SMNs (Figure 2a, 2b). Regression analysis using Lm model was performed to determine correlation between the gene expression changes in the two datasets.

#### Expression of gene modules

AddModuleScore was used to plot average expression of gene modules on UMAP such as in Figures 3a, 3b, 3c, 4a, 4b, 4c or average accessibility at open chromatin regions such as in Supplementary Figure 3. This was also used to determine the percent of cells expressing a specific module over a threshold in Figure. 3d. Gene and chromatin modules that define each cell type was determined by using FindMarkers.

#### Label Transfer Analysis

All label transfer analysis was performed using the RNA-seq data. Seurat Objects from all ages containing just SMNs (or PGMNs) were independently clustered. Clusters at E15.5 and P21 were assigned identities based on known pool/column genes and alpha, gamma, type3 markers respectively. Then, FindTransferAnchors (normalization.method = "SCT", reference.reduction = "pca", dims = 1:50) and MapQuery(reference.reduction = "pca", reduction.model = "umap.rna") were used to assign reference identities to query datasets. In Figure 3, E15.5 identities were the reference identities, while all other ages were used as query one at time. In Figure 4, P21 identities were the reference identities while all other ages were used as query one at a time. In this analysis, each query nucleus is given a predicted score ranging from 0 to 1 for each reference cluster. If a nucleus has a predicted score of 0.5 for two reference clusters, this suggests that it is equally similar to both clusters; for this reason we only assigned a reference cluster identity to a query nucleus if it has a predicted score > 0.6. All nuclei with a predicted score < 0.6 were given non-significant match status.

#### Label Transfer Analysis of Data from Alkaslasi et al.^20^

Cervical_MF and Lumbar_MF data was acquired from GSE167597. Matrix, features, and barcode files were converted into Seurat objects and processed as follows: Alkaslasi.p56 <- merge(cervical, lumbar, add.cell.ids = c("cervical", "lumbar"), project = "AlkaslasiP56") Alkaslasi.p56 <- SCTransform(Alkaslasi.p56, vst.flavor = "v2) %>%

RunPCA(npcs = 30, verbose = FALSE) %>%

RunUMAP(reduction = "pca", dims = 1:30) %>%

FindNeighbors(reduction = "pca", dims = 1:30) %>%

FindClusters(resolution = 0.7)

SMN clusters were identified using the same methods metioned above, namely by identifying clusters that were Chat+ and expressed Slc5a7, Bcl6, Ret, Gfra1, and Tns1. SMN clusters were subset and used for Label Transfer analysis as described above.

#### Determination of E15.5 pool/column markers

Top pool markers were identified by performing FindAllMarkers on E15.5 only SMN clusters from Supplementary Figure 2. Top column markers were identified by performing FindMarkers, comparing LMC, HMC, or MMC clusters to all others.

#### Determination of top alpha, gamma, and type3 markers

FindMarkers was used to find mature alpha (ident.1 = c(9), ident.2 = c(0, 32, 11, 4, 25)), gamma (ident.1 = c(11, 4, 25), ident.2 = c(9,0,32)), and type3 markers (ident.1 = c(32,0), ident.2 = c(9, 11, 4, 25)) from merged dataset of all SMNs shown in Figure 2B. Any genes that showed up as markers for multiple subtypes were eliminated. To identify markers that are even more specific, we used more stringent criteria of (avg_log2FC >=1.4, (pct.1 - pct.2) > 0.2, p_val_adj < 0.001). This left us with 143 top alpha markers, 100 top gamma markers, and 54 top type3 markers. All differential genes and filtered top markers are listed in Supplementary Table 1.

#### Differential ATAC-seq peaks

Column/pool specific peaks, and alpha/gamma/type3 specific peaks were identified by performing FindMarkers on the combined all ages UMAP from Figure 2B. To identify pool specific peaks, all the E15.5 clusters were compared to each other (and not to clusters from other ages). For alpha, gamma, type3 specific peaks comparisons were as follows: alpha (ident.1 = c(9), ident.2 = c(0, 32, 11, 4, 25)), gamma (ident.1 = c(11, 4, 25), ident.2 = c(9,0,32)), and type3 (ident.1 = c(32,0), ident.2 = c(9, 11, 4, 25)).

#### Motif Enrichment

Motif enrichment in differential alpha, gamma, and type3 peaks was performed by using AddMotifs and FindMotifs in Signac. The entire Jaspar2020 motif database was used for this analysis. For visualization of motifs on UMAP, ChromVAR was used to add a motif assay to Seurat objects, human motifs from the Jaspar2020 database were used for this analysis.

### Enhancer AAVs

#### Cloning enhancer-AAVs

We selected 3 differential alpha and gamma open chromatin peaks that were near alpha and gamma specific genes respectively. These regions were amplified out of genomic DNA of C57BL/6J (JAX stock#000664) mice, and cloned into addgene plasmid #193741, which is a previously validated ^29^ <u>AAV</u>- CHAT<u>enhancer</u>-miniProm-chimericIntron-<u>mCherry</u>-WPRE plasmid. The chat enhancer was replaced with new open chromatin peaks by digesting this plasmid with Kpn1 and Age1, and inserting new peaks with Gibson assembly (NEB HiFi Assembly Mix). Chromosome locations of peaks and primers are below. These primers were used to amplify peaks from genomic DNA and simultaneously add homology regions to plasmid backbone for Gibson Assembly.

Gamma 1: chr6-53562382-53563316

Fwd: gtagccatgctctaggaagagtaggtaccAACTCCCTCTGAATGACAACC

Rev: AGCTTCCATTATATACCCTCTAgaccggtCTAGCATCTCTTTCTCTCTCT

Gamma 2: chr12-81175030-81175991

Fwd: gtagccatgctctaggaagagtaggtaccGCTAAAGCTTTGTTACTTAAG

Rev: AGCTTCCATTATATACCCTCTAgaccggtAAGGCACACTGAGCTGTCACT

Gamma 3: 61997587-61998537 (we followed up on this one)

Fwd: agccatgctctaggaagagtaggtaccAATACATAGATGCAGTCTATCAG

Rev: AGCTTCCATTATATACCCTCTAgaccggtCCATTCCAGTCAAATAGAACA

Alpha 1: chr18-43252300-43253300

Fwd: gtagccatgctctaggaagagtaggtaccGATAAACAATGTTTTAGTAGA

Rev: AGCTTCCATTATATACCCTCTAgaccggtTTTCTTTCTTTCTTTTGTATC

Alpha 2: chr8-116030939-116031598

Fwd: gtagccatgctctaggaagagtaggtaccAAGAGCGTGAATCGACTCTTA

Rev: AGCTTCCATTATATACCCTCTAgaccggtCATTTGTCATTAAGATGCACA

Alpha 3: chr11-118546009-118547095 (we followed up on this one)

Fwd: gtagccatgctctaggaagagtaggtaccATGACTTTGAAGCCATACACG

Rev: AGCTTCCATTATATACCCTCTAgaccggtGTTTTGGGCCTCAGCACACAC

#### Generating AAVs

AAVs were generated following the protocol from Negrini et al.^58^ without any changes to the published protocol. AAV titration was performed following the titration protocol in Challis et al.^59^. The PHP.eB cap plasmid^60^ and pHelper along with AAV-enhancer-mCherry constructs to independently generate 6 different viruses, which were independently injected and characterized. AAVs were injected by retroorbital injection at a titer of 3e13 viral genomes per mouse. Injections were performed in P21-P35 mice, and tissues was collected 14 days after injection.

### RNA FISH

#### Probes and Protocol

HCR v3 probes and hairpins from Molecular Instruments were used for multi-color RNA FISH. Probes for all genes were designed by Molecular Instruments. The Molecular Instruments RNA FISH protocol for Frozen Tissue, Rev 4 did not produce good results, but a modification of this protocol as described in Frank et al.^61^ worked well and was used without any further changes. Images were taken at 20x using a Zeiss Axio Observer 7 with Apotome 3. Z-sections were collected every 1um.

#### Tissue preparation

Deeply anesthetized mice were transcardially perfused with 20 ml of PBS, followed by 20 ml of 4% paraformaldehyde. Vertebral columns were dissected out and fixed overnight in 4% paraformaldehyde at 4 °C, followed by storage in PBS. Brachial and Lumbar sections of the spinal cord were dissected, immersed sequentially in 10%, 20%, and 30% sucrose, and then frozen in OCT. Tissue was sectioned at 15-20um, dried overnight at room temperature and stored at −80C.

#### Quantification

Quantification of FISH signal was performed manually in ImageJ, by first counting all cells that expressed Chat, and then counting Chat+ cells that also showed expression of subtype markers of interest. As Chat RNA filled up the whole cell, this was used to define the boundaries of the cell, and consistent FISH signal within those boundaries was counted as a positive signal.

### Immunostaining

#### Tissue preparation and protocol

Deeply anesthetized mice were transcardially perfused with 20 ml of PBS, followed by 20 ml of 4% paraformaldehyde. Vertebral columns were dissected out and fixed for 2 hours in 4% paraformaldehyde at 4 °C, followed by storage in PBS for at least 24 hours. Brachial and lumbar spinal cords were dissected out of the vertebral columns and sectioned with a vibratome in 30–70 μm sections.

Immunostaining was performed in free-floating sections. Sections were incubated in primary antibody for 48 h at room temperature in Antibody Buffer (1% BSA, 0.5% Triton X-100 in 1X PBS, and 0.05% NaN3), followed by 3x 1–3 h washes with Wash Buffer (0.1% Triton X-100 in PBS), followed by secondary antibody treatment in Wash Buffer overnight at room temperature, followed by 1x wash with Wash Buffer + DAPI for 1 h, and 2x washes with just Wash Buffer for 30 min to 1 h each. Sections were then placed on slides and sealed with Flouromount G (Thermo Fisher Scientific OB100-01) and coverslips. Images were taken at 20x using a Zeiss Axio Observer 7 with Apotome 3. Z-sections were collected every 2um.

#### Antibodies

Primary antibodies used were: Chat (Goat, EMD Millipore AB144P 1:100), NeuN (Rabbit, Millipore-Sigma ABN78, 1:1000), Nfia (Rb, Active Motif 39397, 1:1000), Nfib (Rb, Active Motif 39091, 1:1000), Creb5 (Rb, PA5-65593, Thermo Fisher, 1:100), Esrrg (Ms, R&D Systems PPH681200, 1:300), and mCherry (Rt, Thermo Fisher M11217, 1:1000).

Alexa Flour and Jackson Immunolabs secondary antibodies were used at 1:2000 and 1:800 respectively.

#### Quantification

Quantification of enhancer AAVs was performed manually by counting the number of mCherry positive cells that overlapped with Chat and Neun/Esrrg. Quantification of transcription factors staining was performed automatically by ImageJ. Nuclei were circled as regions of interest and mean intensities of Esrrg and TF channels within these regions of interest were determined by ImageJ. The mean intensities for Esrrg and TFs were independently normalized ((value-min)/(max-min)) within each image to control for staining variability between sections. The normalized values of Esrrg and TF intensities were plotted in Figure 6.

Pearson correlation between Esrrg and TF values were calculated.

## Acknowledgements

The authors would like to thank Hynek Wichterle, Esteban Mazzoni, Vilaiwan Fernandes, and Oliver Hobert for critical feedback on the manuscript and scientific discussion. We would also like to thank Michael Kissner and the Columbia Stem Cell Initiative Flow cytometry core, as well as the Columbia Genome Center at Columbia University. This work was supported by the NIH (K99-NS121136, R00NS121136), Project ALS, and startup funds from Rutgers Robert Wood Johnson Medical School to TP.

## Author contributions

T.P. conceived the project, performed snMultiome experiments, analyzed the snMultiome data, helped design and supervise all experiments, and wrote the first draft of the manuscript. Y.C. designed, performed, and analyzed the AAV experiments, and edited the manuscript. H.C performed and analyzed the RNA FISH experiments, and helped with AAV experiments. A.T. performed and analyzed transcription factor immunostainings, and edited the manuscript. A.M. helped with RNA FISH experiments and edited the manuscript.

## Materials Availability

Plasmids generated in this study are available from the lead contact with a complete materials transfer agreement.

## Data and code availability

- Single nucleus multiome data will be made publicly available at GEO upon publication in a peer-reviewed journal.
- Microscopy data reported in this paper will be shared by the lead contact upon request.
- All code will be made available on Github upon publication in a peer-reviewed journal.
- Any additional information required to reanalyze the data reported in this paper will be available from the lead contact upon request.

## Lead Contact

Requests for further information and resources should be directed to and will be fulfilled by the lead contact, Tulsi Patel

## Figure Legends

**Supplementary Figure 1:**
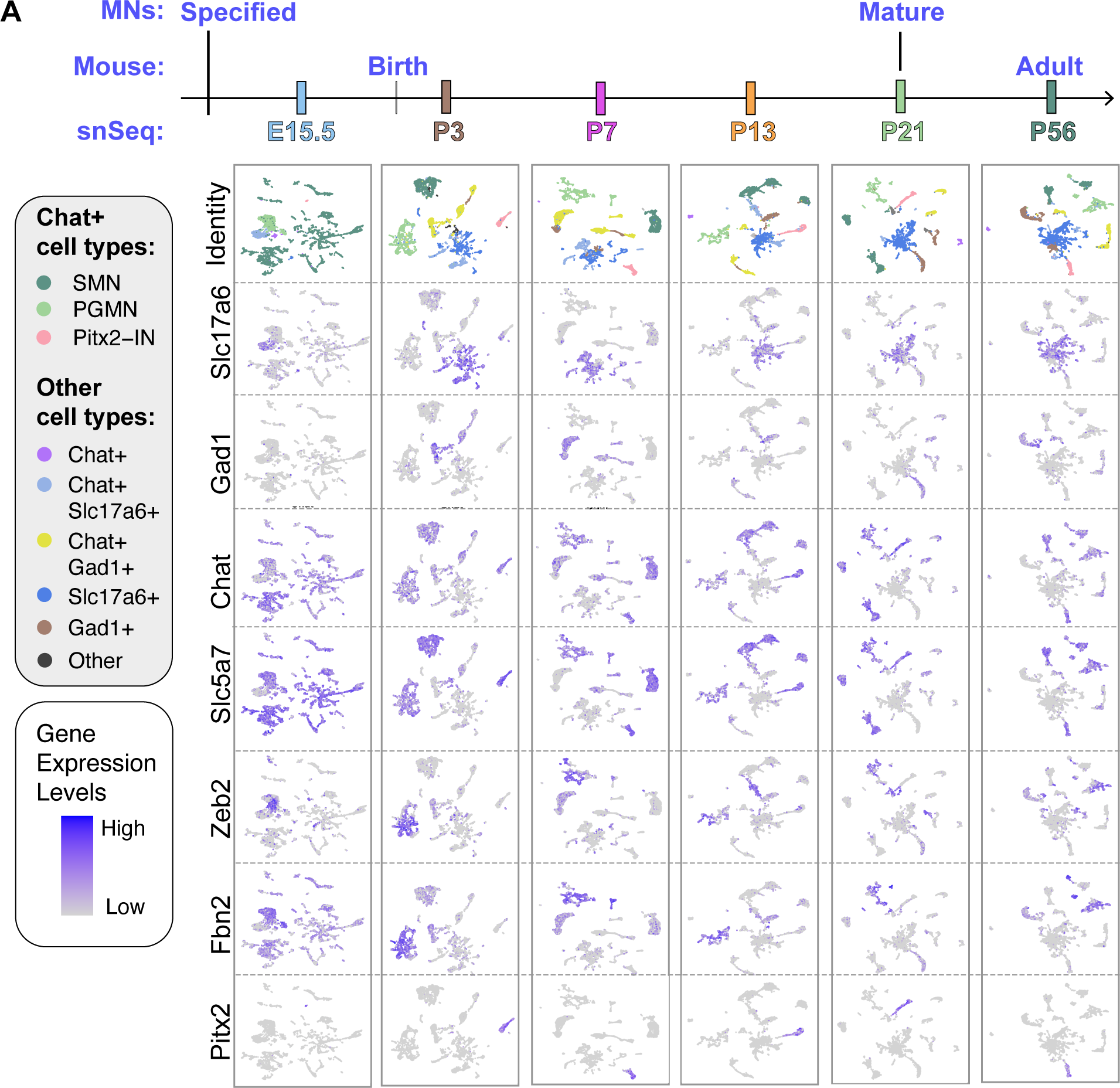
Marker expression in single nucleus multiome trajectory. **A)** Extension of Figure 1a. UMAP representation of all nuclei that passed quality control at each age. Identity of nuclei is determined by expression of cholinergic genes Chat and Slc5a7, GABAergic gene Gad1, and excitatory gene Slc17a6, as well as cell type specific markers including Zeb2 and Fbn2 for PGMNs.

**Supplementary Figure 2:**
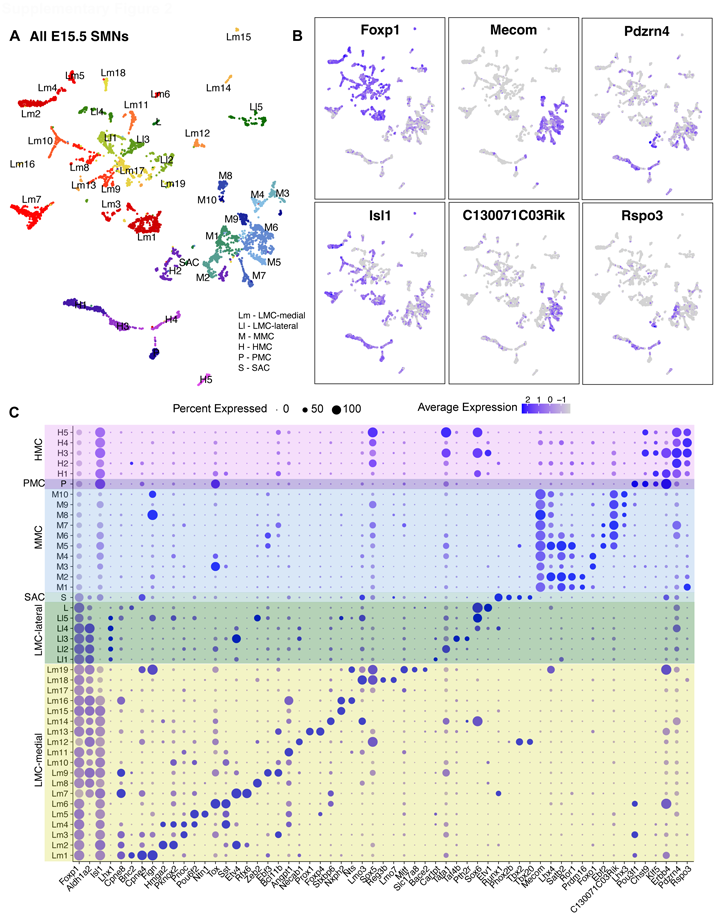
Analysis of column and pool diversity in E15.5 SMNs. **A)** Clustering of all SMNs from E15.5. Column identities LMC (lateral motor column), MMC (medial motor column), HMC (hypaxial motor column), PMC (phrenic motor column) and SAC (spinal accessory column) are assigned based on gene expression as demonstrated in **B** and **C**. LMC can be further divided into lateral and medial based on expression of Isl1 or Lhx1.

**Supplementary Figure 3:**
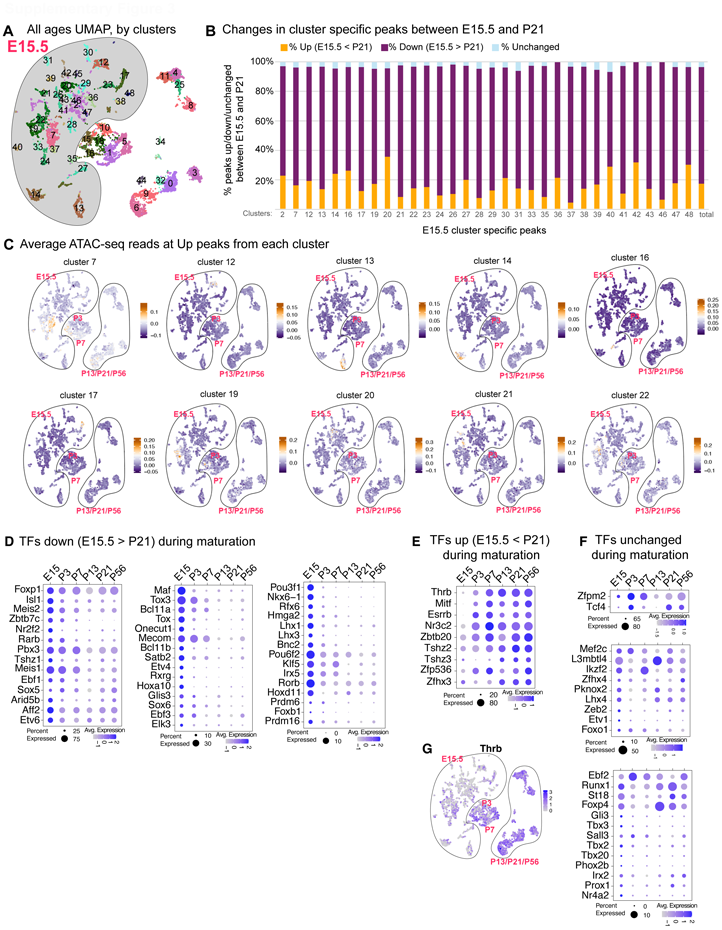
Chromatin accessibility and TF profiles of E15.5 column/pool diversity are lost over time. **A)** All ages UMAP from Figure 2b, with E15.5 clusters highlighted. **B)** For each E15.5 cluster in (A), percent of cluster-specific peaks that show increased accessibility (Up), decreased accessibility (Down), or unchanged accessibility over time. **C)** Average accessibility at cluster-specific peaks that are Up in SMNs of all ages. **D, E, F**) Expression of TFs that that show cluster specific expression in E15.5 clusters. TFs that are significantly downregulated between E15.5 and P21 are in (D), significantly upregulated in (E) and unchanged in (F). **G)** Expression of Thrb, which is upregulated over time, in all SMNs.

**Supplementary Figure 4:**
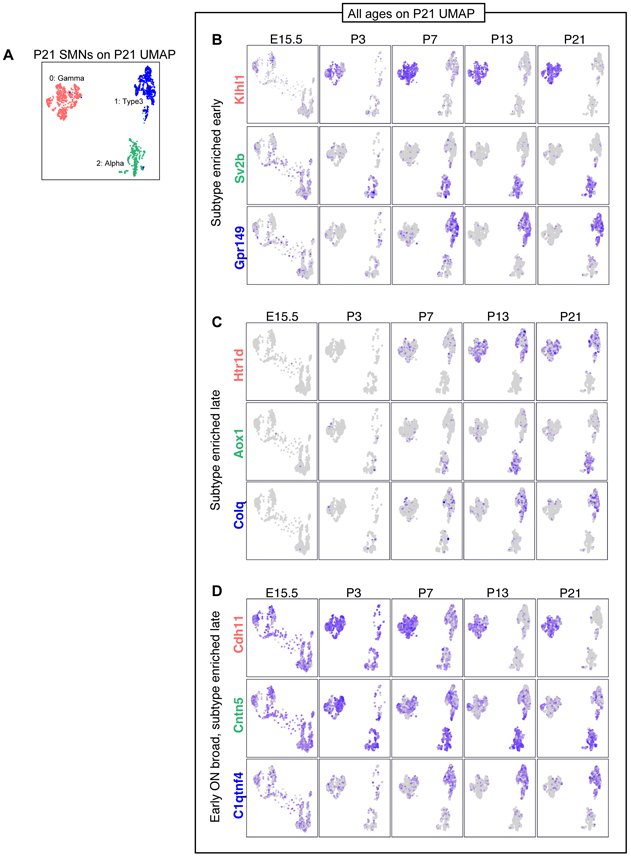
Alpha, Gamma, and Type3 subtype-specific genes are sequentially enriched during maturation. **A)** UMAP of all P21 SMNs. **B, C, D)** E15.5, P3, P7, P13 SMNs plotted on P21 UMAPs based on label transfer analysis. Examples of different categories of alpha, gamma, and type3 subtype genes are shown. Genes that show subtype specific expression early are in (B), genes that show subtype specific expression later are in (C), and genes that are broadly active early but become more specific later are in (D). Early and late is relative for each subtype as each subtype acquires subtype specific gene expression patterns over different time scales.

